# *mucG, mucH*, and *mucI* modulate production of mutanocyclin and reutericyclins in *Streptococcus mutans* B04Sm5

**DOI:** 10.1101/2022.01.27.478122

**Authors:** Jonathon L. Baker, Xiaoyu Tang, Sandra LaBonte, Carla Uranga, Anna Edlund

## Abstract

*Streptococcus mutans* is considered a primary etiologic agent of dental caries, which is the most common chronic infectious disease worldwide. *S. mutans* B04Sm5 was recently shown to produce reutericyclins and mutanocyclin through the *muc* biosynthetic gene cluster, and to utilize reutericyclins to inhibit the growth of neighboring commensal Streptococci. In this study, examination of *S. mutans* and *muc* phylogeny suggested evolution of an ancestral *S. mutans muc* into three lineages within one *S. mutans* clade, and then horizontal transfer of *muc* to other *S. mutans* clades. The roles of the *mucG* and *mucH* transcriptional regulators, and the *mucI* transporter, were also examined. *mucH* was demonstrated to encode a transcriptional activator of *muc. mucH* deletion reduced production of mutanocyclin and reutericyclins, and eliminated the impaired growth and inhibition of neighboring Streptococci phenotypes which are associated with reutericyclin production. Δ*mucG* had increased mutanocyclin and reutericyclin production, which impaired growth and increased the ability to inhibit neighboring Streptococci. However, deletion of *mucG* also caused reduced expression of *mucD, mucE*, and *mucI*. Deletion of *mucI* reduced mutanocyclin and reutericylin production, but enhanced growth, suggesting that *mucI* may not transport reutericyclin as its homolog does in *Limosilactobacillus reuteri*. Further research is needed to determine the roles of *mucG* and *mucI*, and to identify any co-factors affecting the activity of the *mucG* and *mucH* regulators. Overall, this study provided pangenome and phylogenetic analysis that serves as a resource for *S. mutans* research, and began elucidation of the regulation of reutericyclins and mutanocyclin production in *S. mutans*.

**Importance:** *S. mutans* must be able to outcompete neighboring organisms in its ecological niche in order to cause dental caries. *S. mutans* B04Sm5 inhibited the growth of neighboring commensal Streptococci through production of reutericyclins via the *muc* biosynthetic gene cluster. In this study, carriage of *muc* was examined across *S. mutans*, which showed that 35 of 244 RefSeq *S. mutans* genomes encoded *muc* and provided a valuable update to the *S. mutans* pangenome and phylogeny. The roles of the *mucG* and *mucH* transcriptional regulators, and the *mucI* transporter, were also examined. All 3 genes impacted production of mutanocyclin and reutericyclins, which affected the growth rates, transcriptomes, and the ability of the *S. mutans* strains to inhibit the growth of neighboring commensals.

## Introduction

*Streptococcus mutans* is considered a primary etiologic agent of dental caries, which is the most common chronic infectious disease worldwide (1). As it is not typically considered a pioneer colonizer of the tooth, *S. mutans* must be able to out-compete already established bacterial neighbors (which are typically health-associated commensals) to successfully establish itself as a member of the dental plaque microbiota and cause disease (2, 3). To achieve this outcome, *S. mutans* uses several different abilities. *S. mutans* is able to generate insoluble glucans from sucrose, which greatly facilitate biofilm formation on the tooth surface (4). Therefore, in the presence of a carbohydrate-rich diet by the human host (particularly one with frequent consumption of sucrose), *S. mutans* is at a distinct advantage compared to many of its more health-associated neighbors (2, 3). In addition, *S. mutans* utilizes these host dietary carbohydrates in energy metabolism. This process generates organic acids which quickly and drastically lower the local pH, which is what causes damage to the underlying tooth (1). While *S. mutans* employs a complex and robust acid tolerance response to continue to thrive in these acidic conditions, many of its health-associated competitors are much more acid sensitive and cannot sustain growth (5). In addition to these somewhat indirect competition strategies, *S. mutans* also directly inhibits the growth of its competitors through the production of antimicrobial small molecules, such as bacteriocins (which are termed mutacins in *S. mutans*) (6).

One group of these antimicrobial small molecules are reutericyclins, which are acylated tetramic acids produced by a biosynthetic gene cluster (BGC), *muc* (Figure S1), encoded by a subset of globally distributed *S. mutans* strains (7-9). The *muc* BGC consists of 10 genes (*mucA-J*) (8, 9). *mucD* and *mucE* encode the biosynthetic core genes; a nonribosomal peptide synthetase (NRPS) and a polyketide synthase (PKS), respectively. *mucA, mucB*, and *mucC* are predicted to encode tailoring enzymes (8). *mucF* was recently shown to encode a previously unknown class of acylase with an HXXEE motif, while *mucG* and *mucH* are predicted to encode TetR/AcrR family transcription regulators. *mucI* is predicted to encode a DHA2 family transporter, and *mucJ* is predicted to encode a small multi-drug export protein (8, 9). In *S. mutans* B04Sm5, *muc* produces four tetramic acid compounds; three reutericyclin molecules (differing in the length and saturation of the acyl chain, and referred to collectively in this manuscript as simply “reutericyclin”) and the unacylated tetramic acid, mutanocyclin (8). Previous experiments illustrated that deletion of the gene encoding the MucF acylase abolishes production of unacylated mutanocyclin (8). As a result, the Δ*mucF* strain accumulates more reutericyclin molecules, which leads to growth inhibition of both itself and neighboring health-associated oral Streptococci (8). Recently, reutericyclin treatment was shown to inhibit biofilm formation and acid production by an *in vitro* oral microbiome community, as well as significantly alter the taxonomic profile of the community (10). Meanwhile, mutanocyclin did not have any antimicrobial activity against several species of oral Streptococci, but demonstrated anti-inflammatory activity in a murine model (9). Mutanocyclin treatment did not alter the taxonomic profile of an *in vitro* oral biofilm community to the same extent as reutericyclin, but did significantly reduce the abundance of *Limosilactobacillus fermentum*, specifically (10). Other possible roles of mutanocyclin on *S. mutans* metabolism and ecology, including if mutanocyclin has direct bactericidal or bacteriostatic activity against *L. fermentum*, are the subject of current investigation. As the majority of research on *S. mutans* has been conducted using well established type strains which do not encode *muc* (such as UA159 and UA140) the roles of *muc*, and its products within the *S. mutans* lifestyle are not well understood.

In this study, the complete genome of B04Sm5, which was recently reported (11), was analyzed, and pangenome analysis of 244 *S. mutans* genomes from NCBI was performed to examine the distribution of the genes within the *muc* BGC. Additionally, the roles of *mucG, mucH*, and *mucI* vis-a-vis the production of mutanocyclin and reutericyclin, regulation of *muc*, overall transcriptome, and ability of *S. mutans* to inhibit neighboring commensals were further investigated.

## Results

### Comparative genomics of B04Sm5, the *S. mutans* pangenome, and distribution of *muc*

The complete genome sequence of *S. mutans* B04Sm5, the strain where reutericyclin and mutanocyclin production was recently characterized, has now been reported (11). Compared to the *S. mutans* type strain, UA159, B04Sm5 has a large ∼1.4Mbp X-shaped chromosomal inversion, as did NN2025, another *S. mutans* isolate with a published complete genome sequence (12), which also encodes *muc* (Figure 1A). To examine the distribution of the genes of the *muc* BGC across the *S. mutans* pangenome, the genomes of 244 *S. mutans* strains from GenBank (all *S. mutans* genomes on GenBank, as of June 2021; Table S1) were examined using Anvi’o (13). Using the parameters described in the Materials and Methods section, the *S. mutans* pangenome included 3,183 genes. The pangenome data table file was too large to be included as Supplemental Material, but is publicly available at https://github.com/jonbakerlab/Smutans-pangenome. There were 1,421 genes in the core genome (present in >90%, or 232, of the genomes), 1,212 genes in the cloud pangenome (present in <10%, or 36 of the genomes), and 549 genes in the shell pangenome (present in ≥10% and ≤90% of the genomes) (Figure 1B). *muc* was found on the boundary from the cloud to the shell pangenome, present in 35 of the 244 genomes, with all genes in the BGC (Anvi’o gene calling did not identify *mucJ*, but a subsequent manual search did) present in all 35 of these strains, indicating there were no versions of the *muc* BGC in *S. mutans* missing any of the 10 genes (https://github.com/jonbakerlab/Smutans-pangenome). Concatenated protein sequences of MucA-I were used to construct a phylogenetic tree of the *muc* BGC within *S. mutans*, which demonstrated three main lineages of Muc (Figure 1C). To search for other genes that may affect whether it is advantageous for *S. mutans* to encode *muc*, the pangenome was examined for genes that co-occurred with the *muc* BGC using Coinfinder (14). Forty-four genes had their carriage correlated to that of *muc* (Figures 1D, S2, Table S2). Twenty-eight of the 44 genes associated with *muc* were found in B04Sm5, however none were located adjacent *muc* in the chromosome. Notable among the correlated genes were another hybrid NRPS/PKS BGC (GC00001910) and a lantipeptide BGC (GC00001893). No genes were significantly negatively correlated with carriage of *muc*, using the parameters employed, indicating that there may be no genes in the *S. mutans* pangenome that contraindicate carriage of *muc* or the production of reutericyclin or mutanocyclin.

**Figure 1:**
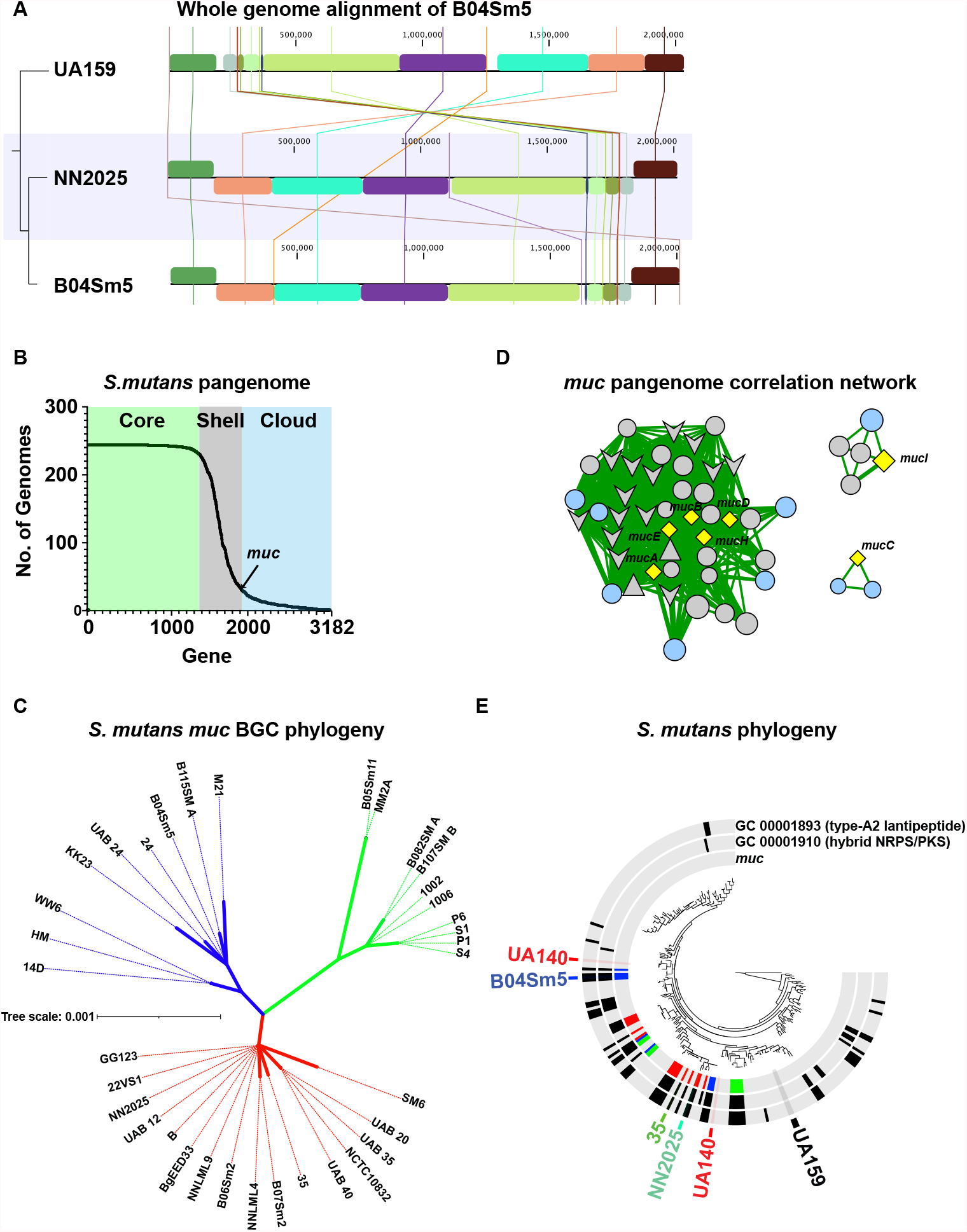
Comparative genomics of B04Sm5 and the pangenome of *S. mutans*. **(A)** Whole genome alignment of *S. mutans* B04Sm5 versus the type strains UA159 (- *muc*) and NN2025 (+ *muc*). The tree on the left is based on the whole genome alignment itself. **(B)** The *S. mutans* pangenome. Graph showing the gene clusters in the *S. mutans* pangenome versus the number of genomes that each gene appears in. The Core (>90% of genomes), Shell (≥10% and ≤90% of genomes), and Cloud (<10% of genomes) pangenomes are indicated by the background colors green, gray, and blue, respectively. The location of *muc* on the graph is indicated by the arrow. **(C)** *S. mutans muc* BGC phylogeny. Concatenated protein sequences of MucA-I across the 35 *S. mutans* genomes harboring *muc* were used to construct an unrooted phylogenetic tree of the gene cluster. The three main lineages are colored blue, green, and red. **(D)** *muc* pangenome correlation network. Correlation network illustrating the *muc* genes (yellow diamond nodes) and genes they are correlated with, based on Coinfinder analysis. Node color of non-*muc* genes indicates presence in the shell (gray) or cloud (blue) pangenome. Node shape indicates gene clusters of interest: diamond = *muc*; chevron = hybrid NRPS/PKS; triangle = type-A2 lanthipepetide; circle = all other genes. Edges represent positive correlations only, and edge thickness indicates the p-value of the correlation. The correlation network, with all nodes labeled is available in Figure S2. **(E)** *S. mutans* phylogeny. Phylogenetic tree of 244 *S. mutans* genomes based on the concatentated protein sequences of 12 core genes, as described in the Materials and Methods. The tree is annotated with 3 layers indicating the presence of *muc*, GC0000189 (type-A2 lanthipeptide which was correlated with muc in Panel D), and GC00001910 (hybrid NRPS/PKS which was correlated with *muc* in Panel D). Presence/absence bars in the *muc* layer are colored based on the position of the *muc* BGC of the cognate genome in the *muc* phylogenetic tree in Panel C. *S. mutans* genomes of interest are labeled (B04Sm5, UA159, UA140, NN20205, and 35). The full phylogenetic tree, with all leaves labeled, is available in Figure S3.

To further examine phylogeny of *S. mutans*, and distribution of *muc*, the pangenome was used to identify optimal protein sequences to use in constructing a phylogenetic tree of the 244 *S. mutans* strains. The pangenomics analysis identified 348 single-copy core genes encoded by all 244 genomes. Using an approach described by (13), these were filtered by a minimum geometric homogeneity index of one to remove protein sequences that would introduce gaps in the alignment, which left 270 protein sequences. Many of these genes had nearly identical sequences, therefore a further filter of maximum functional homogeneity index was set to 0.9925, leaving 12 genes (listed in Table S3) on which to base the phylogenetic analysis (Figures 1E, S3). The strains encoding *muc* were not monophyletic, and in some cases the three main clades of *muc* did not line up with overall *S. mutans* phylogeny. These results suggest horizontal transfer of *muc S. mutans* clades, as has been suggested for *Lactobacillus spp*. carrying the closely-related reutericyclin BGC (15).

In terms of arsenal of small molecules produced by BGCs other than *muc*, the B04Sm5 genome encodes the type-A2 lanthipeptide bacteriocin mentioned above (found in 51 genomes across *S. mutans*, not in UA159), mutacin IV / non-lanthibiotic mutacin *nlmAB* (found in 136 *S. mutans* genomes) (16-18), mutacin V / *cipB* (various genes found in 91-243 *S. mutans* genomes) (16, 17), and mutacin VI / *nlmD* (found in 229 *S. mutans* genomes) (19, 20), as well as the *nlmTE* transporter to export these non-lanthibiotic mutacins (present in all 244 *S. mutans* genomes) (21) (Table S3). B04Sm5 does not encode the BGC for mutanobactin (22), which is present in ∼95 genomes, including UA159, or the ribosomally synthesized and posttranslationally modified peptide RaS-RIPP BGC (*cidAB*) (23), which is found in UA159, UA140, and NN2025 (18) (and 181 *S. mutans* genomes in total). These gene clusters of interest are listed in Table S3.

### Deletion of *mucG, mucH*, or *mucI* affects production of mutanocyclin and reutericyclin

To examine the roles of the predicted TetR/AcrR family transcriptional regulators within the *muc* BGC, *mucG* and *mucH*, as well as the DHA2-like family transporter, *mucI*, single gene deletion mutants were generated as described in the Materials and Methods section. To illustrate that any phenotypes observed in the mutant strains were not due to polar effects, complement strains (*mucG*-C, *mucH*-C, and *mucI*-C) were produced where the gene of interest was re-inserted in a distant locus, as described in the Materials and Methods section. The deletion mutants, Δ*mucD* and Δ*mucF*, were described previously (8). Cultures of these strains, as well as Δ*mucD*, Δ*mucF*, and the parent strain B04Sm5 were analyzed by HPLC to observe production of mutanocyclin and the reutericyclins. As seen in the previous study (8), deletion of *mucD* abolished production of all four tetramic acids produced by *muc*, while deletion of *mucF* eliminated production of the unacylated mutanocyclin and increased production of the acylated reutericyclins (Figure 2A). Compared to B04Sm5, deletion of *mucG* increased production of both mutanocyclin and the reutericyclins, while deletion of *mucH* or *mucI* decreased production of both mutanocyclin and the reutericyclins, with Δ*mucH* having lower production than Δ*mucI* (Figure 2A). Complementation with *mucG* and *mucH* reversed the phenotype observed in the cognate mutant, indicating that the phenotypes observed were true effects of the deletion of *mucG* or *mucH*, and not simply polar disruptions in the expression of the adjacent genes (Figure 2A). In fact, the phenotypes were reversed to a greater extent than was observed for B04Sm5 (i.e. *mucG-*C had lower production of the tetramic acids than B04Sm5, while *mucH*-C had higher production than B04Sm5) (Figure 2A). This may indicate that expression of *mucG* and *mucH* may be increased in the complement strains, where transcription is driven by the *gtfA* promoter, compared to transcription driven by the cognate native loci promoters in B04Sm5. Meanwhile, both Δ*mucI* and *mucI-*C had reduced tetramic acid production compared to B04Sm5 (Figure 2A). Taken together, these results suggest that *mucG* is a transcriptional repressor of the *muc* operon, while *mucH* is a transcriptional activator. Since it is supernatants, and not cell lysate, being analyzed here, it is possible that MucI exports reutericyclin and/or mutanocyclin, as reflected by the lower amount detected in the supernatant of the Δ*mucI* strain.

**Figure 2:**
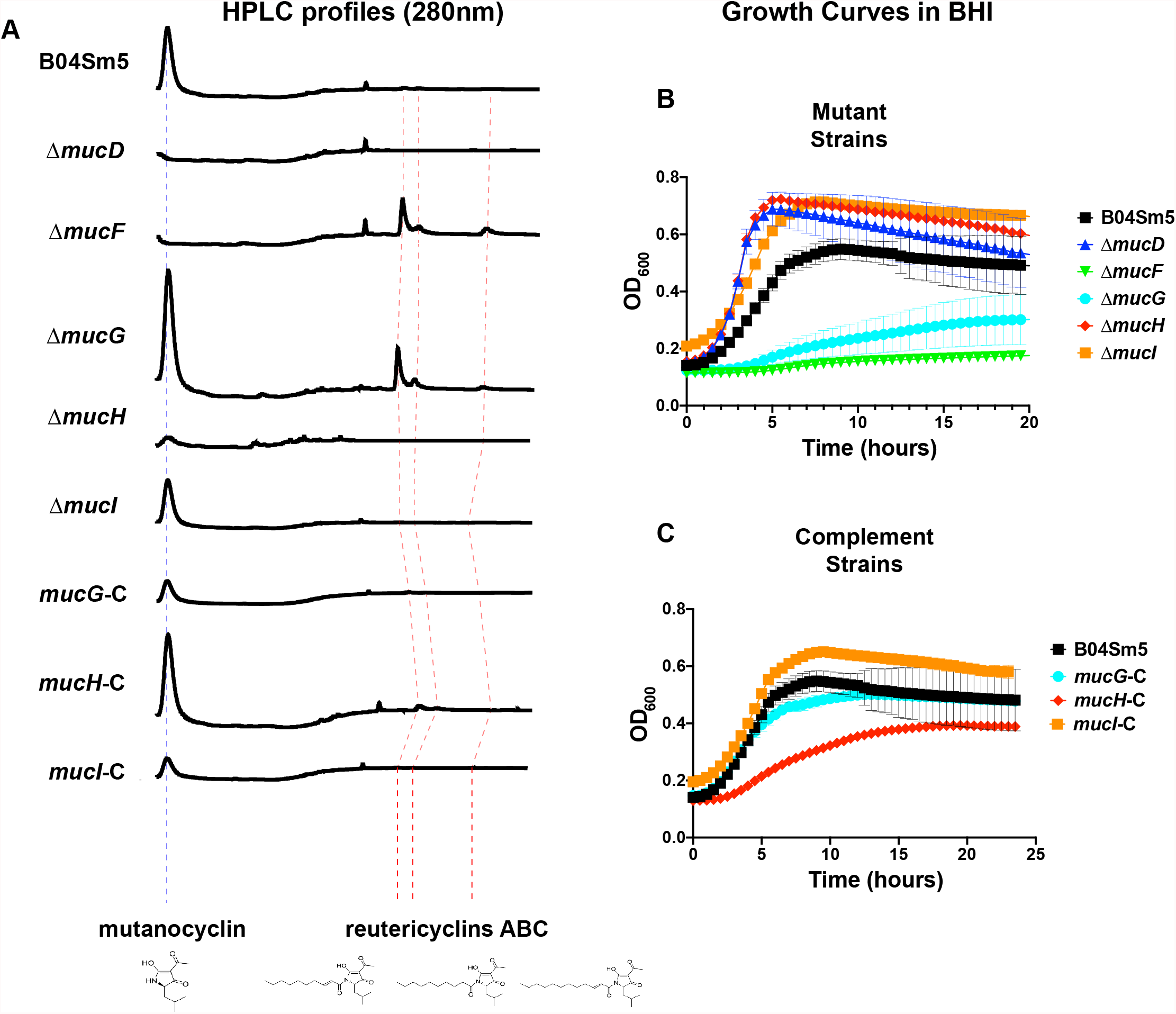
Deletion of the *mucG, mucH, or mucI* impacts the production of mutanocyclin and reutericyclins and the growth of B04Sm5. **(A)** HPLC profiles of concentrated extracts from the supernatant of the indicated strain. Profiles are cropped to retention time (RT) 17-31 min, aligned to the mutanocyclin peak (indicated by the dashed blue line) and presented on the same scale. Red dashed lines indicate the peaks of reutericyclins A B and C. **(B-C)** Growth of the parent strain B04Sm5, and its Δ*mucD, ΔmucF, ΔmucG, ΔmucH*, and Δ*mucI* derivatives (B) or complement strains *mucG*-C, *mucH*-C, and *mucI*-C (C) in BHI. n = 8 for each strain.

### Deletion of *mucG* results in impaired growth while deletion of *mucH* or *mucI* results in improved growth

Growth curves of the Δ*mucD*, Δ*mucF*, Δ*mucG* and Δ*mucH* strains, as well as the complement strains, and the parent strain, B04Sm5, were performed in BHI media. Deletion of *mucG* resulted in significantly impaired growth, nearly to the same level as the Δ*mucF* strain (Figure 2B). Meanwhile, deletion of *mucH* or *mucI* resulted in increased growth, reminiscent of the Δ*mucD* strain (Figure 2B). Complementation of all three genes reversed the phenotypes observed in the cognate mutant strain (Figure 2C). While the *mucG*-C and *mucI*-C strains restored a growth rate similar to the B04Sm5 parent strain, the mucH-C strain actually decreased growth much further, nearly to the level of the Δ*mucG* strain (Figure 2C). Collectively, these results are in line with the HPLC data above, and consistent with the hypothesis that production of the reutericyclins inhibits *S. mutans* growth in a dose-dependent manner.

### Deletion of *mucG* reduces transcription of *mucD, mucE*, and *mucI*, while deletion of *mucH* reduces transcription of the entire *muc* operon

To examine the role of the *mucG* and *mucH* genes that encode transcriptional regulators on the global transcriptome of B04Sm5, mRNA sequencing of mid-log phase cultures of B04Sm5, Δ*mucG*, and Δ*mucH* was performed. Using DESeq2 differential abundance analysis, with a Benjamini-Hochberg corrected p-value cutoff of 0.001, compared to the parental strain, B04Sm5, there were 106 genes with reduced mRNA expression and 218 genes with increased expression in Δ*mucG*, only (Table S4). Meanwhile, compared to B04Sm5, there were 160 genes with reduced expression and 180 genes with increased expression in Δ*mucH*, only (Table S5). There were also 269 genes differentially regulated compared to B04Sm5 in both Δ*mucG* and Δ*mucH*, with all but 11 of those genes changing expression the same direction in both mutants compared to B04Sm5 (Table S6). In terms of expression of the *muc* BGC specifically, in both cases, the deleted gene was the gene with the highest reduction in expression, as expected with deletion mutants (Table 1). The Δ*mucG* strain had modestly reduced (−1.5 to -2 log_2_-fold) expression of *mucD, mucE*, and *mucI*, while the Δ*mucH* strain had more significantly reduced (−2.1 to -8.5 log_2_-fold) expression of the entire operon (Table 1). Given the HPLC and growth phenotypes observed, it was surprising that Δ*mucG* had reduced transcription of *mucD, mucE*, and *mucI*. This suggested that the cause of the increased production of mutanocyclin and the reutericyclins observed in Δ*mucG* must occur at the proteomic or metabolomic level. To obtain an overview of the global transcriptomic effects of deletion of *mucG* or *mucH*, KOs from differentially regulated genes were projected onto a map of *S. mutans* metabolism using KEGG Mapper (https://www.genome.jp/kegg/mapper/color.html; Figure 3). An interactive version of the map can be obtained by the reader by using Table S7 as input for the KEGG Mapper tool. Overall, the results were difficult to interpret and few clear patterns were apparent. The large number of differentially expressed genes was also unexpected, and is likely to reflect changes due to the presence of mutanocyclin and/or reutericyclins, rather than direct regulation by *mucG* or *mucH*. Because Δ*mucG* and Δ*mucH* have opposing phenotypes, the large amount of overlap in the transcriptomes was surprising. It might be expected that Δ*mucH* may reflect the “least stressed” condition, based on the growth curve, and some trends in Δ*mucH* are similar to those seen when *S. mutans* is grown in “non-stressful” conditions (increased expression of *rmlA*), however some trends do not (decreased expression of *accA-D*) (Tinder et al., in review).

**Table 1:** Differential expression of *muc* in the Δ*mucD*, Δ*mucF*, Δ*mucG*, and Δ*mucH* strains, compared to B04Sm5 (log2 fold-change)

| <u>name</u> | <u>locusid</u> | <u>gene_cluster_id</u> | <u>gene_callers</u> | <u>NCBI PGAP ACC</u> | <u>NCBI PGAP</u> | <u>$\Delta mucD$</u> | <u>$\Delta mucF$</u> | <u>$\Delta mucG$</u> | <u>$\Delta mucH$</u> |
| --- | --- | --- | --- | --- | --- | --- | --- | --- | --- |
| *mucA* | IBL27_RS09110 | GC_00001982 | 1693 |  |  | 2.58564569 | 1.19925521 |  | -7.1955871 |
| *mucB* | IBL27_RS09105 | GC_00001981 | 1692 | WP_002277495.1 | thiolase family protein | 1.32475972 | -1.7582417 |  | -4.7312389 |
| *mucC* | IBL27_RS09100 | GC_00001980 | 1691 |  |  |  | -2.3188811 |  | -6.4796598 |
| *mucD* | IBL27_RS09095 | GC_00001984 | 1690 | WP_002308266.1 | non-ribosomal peptide synthetase | -14.06934 | -1.5771313 | -1.8548904 | -6.5515393 |
| *mucE* | IBL27_RS09090 | GC_00001986 | 1689 | WP_002283498.1 | polyketide synthase | -3.5433655 | -2.1506834 | -2.0506685 | -5.9316069 |
| *mucF* | IBL27_RS09085 | GC_00000014 | 1688 | WP_002277499.1 | HXXEE domain-containing protein | -2.7841312 | -5.865363 |  | -2.1839221 |
| *mucG* | IBL27_RS09080 | GC_00001987 | 1687 | WP_002283497.1 | TetR/AcrR family transcriptional regulator | -3.9166794 | -3.1611157 | -8.5142069 | -3.5935604 |
| *mucH* | IBL27_RS09075 | GC_00001983 | 1686 | WP_002277501.1 | TetR/AcrR family transcriptional regulator | 1.06763813 |  |  | -10.557072 |
| *mucI* | IBL27_RS09070 | GC_00001979 | 1685 | WP_002277502.1 | DHA2 family efflux MFS transporter permease subunit | -3.2679608 | -3.2504807 | -1.600364 | -8.524352 |
| *mucJ* | IBL27_RS09065 |  |  |  | small multi-drug export protein | -2.5618472 | -3.3842254 |  | -2.6864964 |

**Figure 3:**
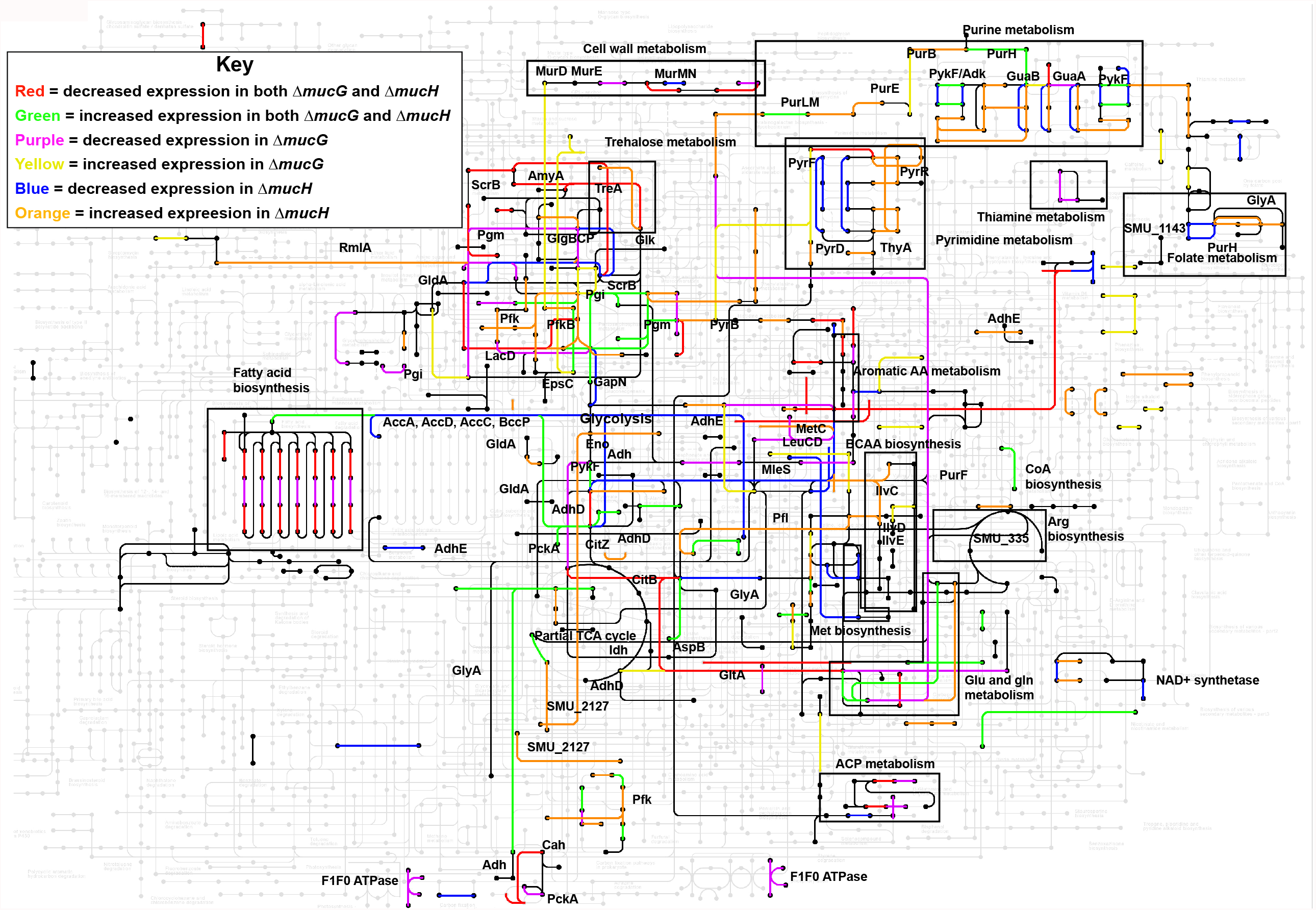
Deletion of *mucG* or *mucH* has broad effects on the B04Sm5 transcriptome. KEGG metabolic map of *S. mutans* metabolism (black nodes and edges) overlaid with the KOs exhibiting differential gene expression deletion of *mucG* (purple for decrease, yellow for increase), *mucH* (blue for decrease, orange for increase), or both (red for decrease, green for increase). Pathways of interest are indicated by labeled boxes, and genes of interest are indicated with labels.

Recently, the transcriptomic effects of addition of wt B04Sm5, Δ*mucD*, or Δ*mucF* to a complex *in vitro* oral microbial community were examined (10). By mapping the mRNA sequencing reads from the complex community in that study to the B04Sm5 genome, a comparison of the transcriptome of Δ*mucD* and Δ*mucF* to Δ*mucG* and Δ*mucH* was possible, however, it is crucial to note that the context of the Δ*mucD* and Δ*mucF* transcriptomes is a community, whereas the Δ*mucG* and Δ*mucH* transcriptomes are single-species cultures (Tables 1, S8, S9). Interestingly, although Δ*mucF* and Δ*mucG*, had similar phenotypes, and Δ*mucD* and Δ*mucH* had similar phenotypes, the overlap between their transcriptomes was small. Altogether, this data indicates that MucH is an activator of transcription of the *muc* BGC, while the role of MucG is less clear.

### Δ*mucG* is significantly better than B04Sm5 at preventing the growth of commensal oral Streptococci, while Δ*mucH* is worse, and Δ*mucI* inhibits only *Streptococcus gordonii*

To examine the ability of the Δ*mucG*, Δ*mucH*, and Δ*mucI* strains in in terms of ability to inhibit the growth of neighboring commensal organisms, colonies of B04Sm5, UA159, the *muc* deletion mutants, and complement strains were spotted on BHI agar or BHI agar buffered to pH 7. Following 24 hours of growth, these plates were overlaid with soft agar containing either *S. gordonii, Streptococcus sanguinis*, or *Streptococcus mitis*, which are all oral Streptococci that are generally health-associated and inversely correlated with *S. mutans* and with dental caries (2, 3). Similar to what was observed in the growth curves described in the growth curves shown in Figure 2, the phenotype of Δ*mucG* mirrored that of Δ*mucF*, while the phenotype of Δ*mucH* mirrored that of Δ*mucD* (Figure 4). Δ*mucF* and Δ*mucG* had significantly larger zones of inhibition against *S. sanguinis* and *S. mitis* compared to the parent strain, B04Sm5. Meanwhile, Δ*mucD* and Δ*mucH* had significantly smaller zones of inhibition against *S. sanguinis* and *S. mitis* compared to B04Sm5 (Figure 4). Buffering the BHI agar to pH 7 reduced the zones of inhibition in most cases, but these phenotypic differences were still observed (Figure 4). This effect was not observed against *S. gordonii*, consistent with a less dramatic effect of *muc* against *S. gordonii* observed previously (8). Interestingly, Δ*mucI* displayed a similar or slightly reduced zone of inhibition against *S. sangunis* and *S. mitis*, but had a significantly increased zone of inhibition against *S. gordonii* at pH 7, but not at pH 5 (Figure 4). As seen in the previous assays, complementation of the Δ*mucG*, Δ*mucH*, or Δ*mucI* mutants faithfully restored the phenotype of the parent strain (Figure 4).

**Figure 4:**
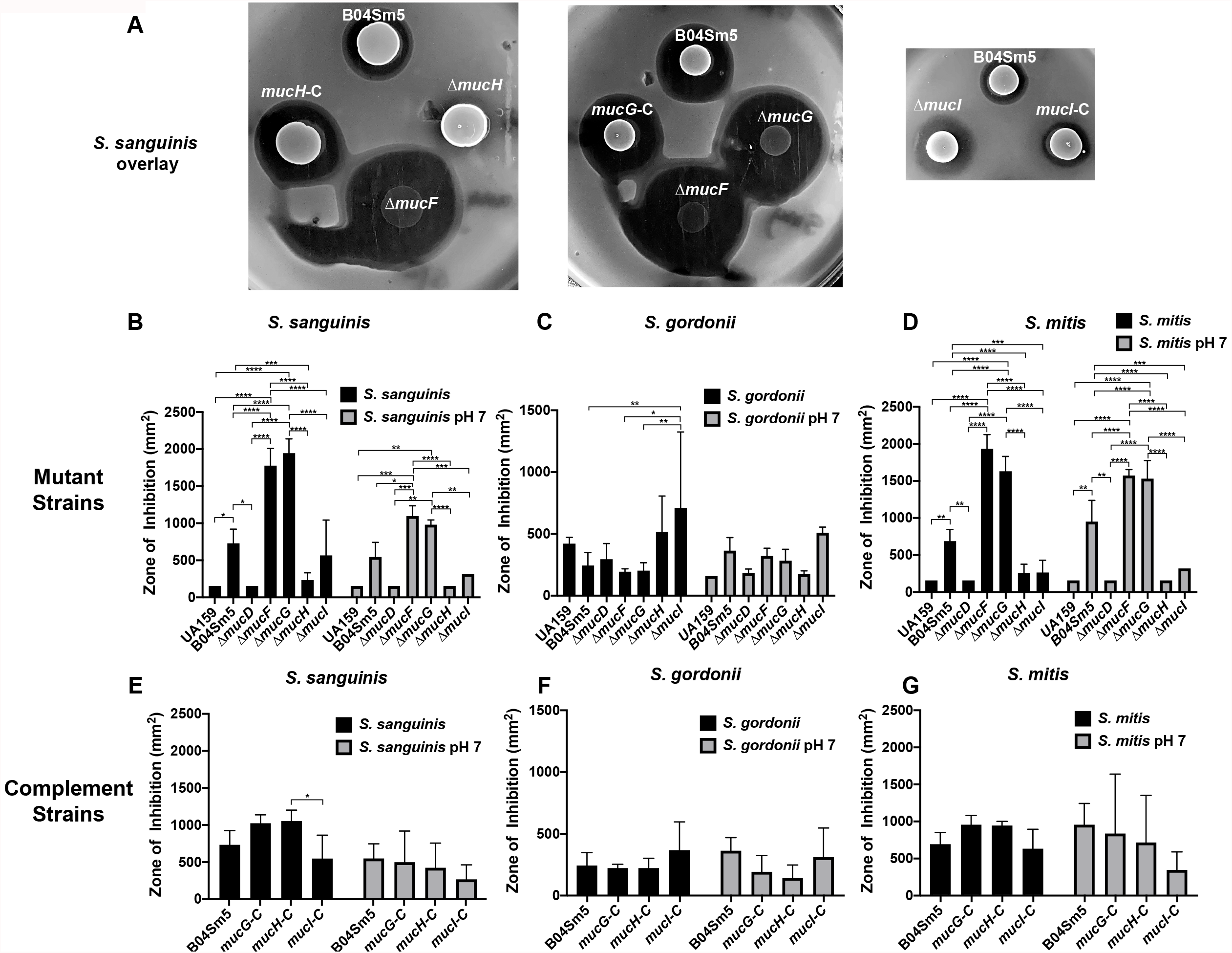
Loss of *mucG, mucH*, or *mucI* impacts the ability of B04Sm5 to inhibit the competing, health-associated bacteria, *S. sanguinis, S. gordonii*, and *S. mitis*. Deferred antagonism assay, performed as described in the Materials and Methods section. Cultures of the indicated *S. mutans* strains were spotted on to BHI agar and incubated overnight. Cultures of the indicated competing organism were added to BHI soft agar and used to overlay the plates containing the *S. mutans*. Three replicate assays were performed, and a representative of the plates with an *S. sanguinis* overlay are shown in panel A. Panels B-G are bar graphs representing quantification of the zones of inhibition observed with the indicated *S. mutans* strain and competing species. Error bars represent standard deviation and asterisks denote statistical significance between indicated pairs as determined by a Tukey’s multiple comparison test following a two-way way analysis of variance (ANOVA)(*, p < 0.05; **, p < 0.01; ***, p < 0.001; ****, p < 0.0001).

## Discussion

Dental caries is caused by a dysbiotic dental plaque microbiome that creates an acidic microenvironment adjacent to the tooth surface, which demineralizes the protective enamel, leading to infection of the dentin and irreparable damage if the process is unchecked (1, 2). *S. mutans* contributes to the formation of this pathogenic community through its exceptional ability to form biofilms in the presence of sucrose, generate acid from many types of dietary carbohydrates, and tolerate an acidic environment (24). Along with indirectly inhibiting health-associated neighbors (which are generally much less acid-tolerant than *S. mutans*) through acid production, *S. mutans* can also directly kill and inhibit the growth of neighboring bacteria through production of small molecules such as class 1 bacteriocins (e.g. mutacin lanthibiotic), non-ribosomal peptides (NRPs), polyketides (PKs) and hybrids of these (e.g. reutericyclin) (8, 24, 25). A large amount of diversity within the *S. mutans* species has been well-documented, both in terms of genomic content, and phenotypes related to virulence traits as mentioned above (26-30). However, it has been difficult to directly link carriages of particular genotypes to dental caries incidence (24, 31, 32).

The updated pangenome provided by this study concurs with previous *S. mutans* pangenomic analyses, indicating that *S. mutans* has approximately 1400 core genes and well over 3000 genes in total (28, 30). Of the 244 *S. mutans* genomes available via the RefSeq database (as of June, 2021), a subset of 35 genomes encode the *muc* BGC. *muc* is homologous to the reutericyclin BGC (Figure S1), which was first elucidated in *L. reuteri*, and produces reutericyclin, an acylated tetramic acid that has potent antimicrobial activity against a broad range of Gram-positive organisms (15). All 35 of the *S. mutans muc* BGCs encode 10 genes: *mucABC*, which currently have unknown function (but are predicted to be tailoring enzymes and are homologous to diacetylphloroglucinol biosynthesis machinery from *Pseudomonas fluorescens*), the MucD NRPS, the MucE PKS, the MucF acylase (which cleaves the acyl chain from reutericyclin to yield mutanocyclin), the TetR/AcrR family transcriptional regulators, MucG and MucH, the MucI DHA2 family transporter, and MucJ, which is predicted to be a small multi-drug export protein. Due to differing versions of antiSMASH software used, MucJ was identified in Hao et al. (9), but not in Liu et al. (7) or Tang et al. (8). Two recent studies independently elucidated that the major product of *S. mutans muc* is mutanocyclin (8, 9), which has the same core tetramic acid structure as reutericyclin, but is lacking the acyl chain. One of these studies also detected production of several types of reutericyclin molecules (differing on the length and saturation of the acyl chain), in addition to mutanocyclin production (8), while the other study did not (9). Although differences in growth conditions and/or biochemical detection methods may account for this, it is also possible that given the sequence divergence of the *muc* cluster between the strains used, that *S. mutans* B04Sm5 produces reutericyclin and *S. mutans* 35 does not (although *mucF*, specifically, is identical between the two strains). Mutanocyclin was anti-inflammatory in a murine model (9) and had limited antimicrobial activity, selectively inhibiting the growth of *Limosilactobacillus fermentum* in a complex *in vitro* oral microbiome (10).

The origin of *muc* remains unclear. In the *L. reuteri* reutericyclin BGC, it was hypothesized that the homologs to *mucA-C* and *mucDE* had distinct phylogenetic origins (15). This was supported by the fact that *rtcN* (*mucD* homolog) and *rtcA* (*mucA* homolog) originated from differing species outside of *S. mutans* and *L. reuteri* (15). The later study by Hao et al. also identified *muc* in *S. troglodytae* and *S. macacae*, which are found in the oral cavities of nonhuman primates (9). Due to modest (40-60% homology) between the *muc* and *rtc* BGCs, direct horizontal gene transfer between *S. mutans* and *L. reuteri* was hypothesized to be unlikely (15). Within *S. mutans*, however, there is little variability in *muc*, with the cluster being 99.4% identical at the protein level, with only 23 sites across *mucA-I* (3,832 amino acids) differing among the 35 strains encoding the gene cluster (https://github.com/jonbakerlab/Smutans-pangenome). Despite these inter-strain differences in *muc* being small, the phylogenetic analysis of *muc* performed here indicated three main lineages of the cluster in *S. mutans* (Figure 1C). These lineages did not line up with overall *S. mutans* phylogeny, suggesting horizontal gene transfer of the cluster within *S. mutans* (Figure 1E). Overall, the data suggests that there may have been one ancestral *muc* within the clade of *S. mutans* containing the most frequent carriage of *muc*, and that this cluster spread to other more distant *S. mutans* clades via horizontal gene transfer. Based on the observations that *muc* is absent in many of the strains within the clade containing the likely ancestral gene cluster also suggests that loss of *muc* is common, and selection pressure may be against retaining *muc* under many conditions. Co-occurrence analysis using Coinfinder (14) identified 46 genes in the *S. mutans* pangenome that co-occurred with *muc* more than would be expected based on phylogeny alone. This indicates that either carriage of *muc* with these genes may provide an advantage over carriage of *muc* or the other genes alone. Many of the co-occurring genes, including the type-A2 lanthipeptide (GC00001893) and the other hybrid NRPS/PKS (GC00001895) are more broadly distributed phylogenetically than *muc* (Figure 1E). Coinfinder also failed to find any genes that contraindicated carriage of *muc*, (i.e. by increasing sensitivity to reutericyclin, for example) however since *muc* is only found in 14% of *S. mutans* genomes and is limited to several clades, it is possible that these types of interactions exist in *S. mutans* and *muc* has simply not sampled enough genetic backgrounds to make these genes apparent statistically. Further examination of the genes and BGCs that correlate with *muc* will determine whether these co-occurring genes increase resistance to reutericyclin or provide synergy with *muc* through another mechanism.

This study also examined the role of the genes encoding the MucG and MucH TetR/AcrR family transcriptional regulators and the MucI DHA2 family transporter. The *L. reuteri rtc* BGC also encodes two TetR/AcrR transcriptional regulators (*rtcRS*), and a transporter (*rtcT*) with varied homology to their counterparts in *S. mutans* (∼20% for *rtcRS/mucGH* and 64% for *rtcT/mucI*) (15). These three genes conferred *L. reuteri* reutericyclin resistance, based on the fact that deletions of *rtcT* and *rtcRS* in reutericyclin-producing strains were lethal, while deletions of these genes in mutants that did not produce reutericyclin were possible (15). Therefore, *rtcT* was predicted to export reutericyclin from the cells, and *rtcR/rtcS* were predicted to be activators of *rtcT* transcription (15). Deletions of the cognate homologous genes, *mucG, mucH*, and *mucI* was accomplished in *S. mutans* B04Sm5, likely because most of the reutericyclin is cleaved into the much less cytotoxic mutanocyclin by *mucF* (8). Future research attempting to delete *mucG, mucH*, or *mucI* in conjunction with *mucF* will test this hypothesis.

Deletion of *mucG* or *mucH* caused opposing phenotypes. Δ*mucG* had increased production of mutanocyclin and reutericyclin, reduced growth, and a larger zone of inhibition of against *S. sanguinis* and *S. mitis*, reminiscent of the phenotype of the Δ*mucF* strain which also produces more reutericyclin (but does not make mutanocyclin, unlike Δ*mucG*)(Figures 2 and 4) (8). Meanwhile, Δ*mucH* mimicked the phenotype of Δ*mucD*, with reduced mutanocyclin and reutericyclin production, enhanced growth, and a significantly reduced zone of inhibition against *S. sanguinis* and *S. gordonii* (Figures 2 and 4) (8). Δ*mucD* does not produce either reutericyclin or mutanocyclin (8). The simple hypothesis from these data would be that *mucG* is a repressor of *muc* and that *mucH* is a transcriptional activator. Based on the transcriptomics, this appears to be the case for *mucH*, as the entire *muc* BGC experienced significantly reduced expression upon its deletion. The role of *mucG* is not as clear, however, as transcription of *muc* was not increased in Δ*mucG*, and in fact expression of *mucD, mucE*, and *mucI* was reduced (however to a lesser extent than was observed in Δ*mucH*). Furthermore, transcription of *mucF* was not decreased, so the ratio of MucD to MucF in Δ*mucG* would appear to be less in favor of producing reutericyclin, which was contrary to the observed phenotype. Since TetR/AcrR regulators typically bind co-factors that affect their transcriptional regulatory activity (33), it is possible that co-factors affect activity of *mucG* and *mucH* and may explain the discrepancy between phenotype and expression level observed in Δ*mucG*. For example, *mucG* may bind either reutericyclin or mutanocyclin and adjust transcription of *muc* to limit reutericyclin production. Alternatively, reutericyclin and mutanocyclin production may be affected in Δ*mucG* by changes at the proteomic or metabolomic level. Additional studies are needed to further elucidate regulation of *muc* by *mucG* and *mucH* and identify the cofactors involved.

The role of MucI remains unclear. The reduced amount of mutanocyclin and reutericlins in the supernatant of Δ*mucI* would seem to indicate that MucI exports one, or both, of these molecules, which is in line with the hypothesis that the *mucI* homolog, *rtcT*, exports reutericyclin in *L. reuteri*. However, deletion of *mucI* also caused enhanced growth, as was seen in Δ*mucD* and Δ*mucH* (Figure 2), and is the opposite of what was observed upon deletion of *rtcT* in *L. reuteri* (15). Accordingly, it is tempting to hypothesize that MucI does not export reutericyclin in *S. mutans*, and instead transports mutanocyclin only, or some other molecule. For example, if MucI transports mutanocyclin, and mutanocyclin is the co-factor that binds either MucG or MucH and turns their transcriptional activation either on or off, that could explain the Δ*mucI* phenotype. It is also possible that MucI somehow regulates and increases production of mutanocyclin and/or reutericyclin. Further research is needed to determine the function of the *mucI* transporter.

Overall, this study provides oral microbiology researchers with an updated phylogenetic analysis of *S. mutans*, and also an updated and in-depth pangenome analysis that can be used to examine carriage of various genes and metabolic functions across *S. mutans* phylogeny. This study also shows that *mucG, mucH*, and *mucI* all affect production of mutanocyclin and reutericyclins by the *muc* BGC and therefore affect *S. mutans* overall physiology. Research in progress will further elucidate the regulation and expression of the *muc* BGC and the production and roles of reutericyclin and mutanocyclin in the ecology and virulence potential of *S. mutans*.

## Materials and Methods

### Bacterial strains and growth conditions

All strains used in this study are listed in Table 2. The *Streptococcus mutans* genomic type strain UA159 (34), B04Sm5 (32), Δ*mucD* (8) and Δ*mucF* (8) strains have been described previously. *S. mutans* was maintained on brain heart infusion (BHI) agar plates (BD/Difco, Franklin Lakes, NJ) at 37°C in a 5% (vol/vol) CO2–95% air environment. Where applicable, antibiotics were added to a final concentration of 5 µg/ml for erythromycin and 1 mg/ml for spectinomycin.

**Table 2:**
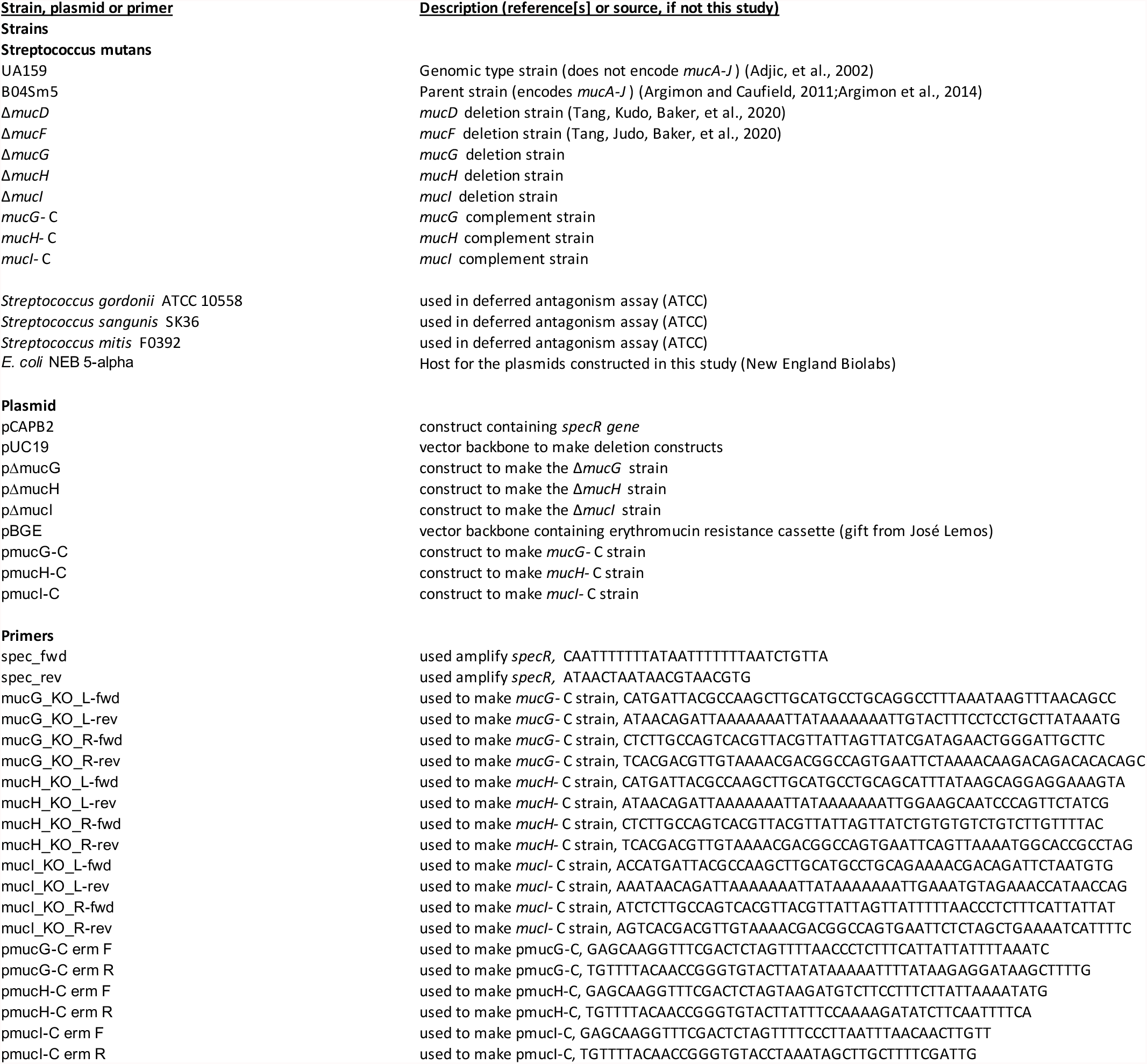
Strains, plasmids, and primers used in this study.

### Pangenome and phylogenetic analyses

Pangenome analysis was performed using the Anvi’o pangenomics Snakemake workflow on 244 *S. mutans* strains from NCBI, listed in Table S1, using DIAMOND, a Minbit parameter of 0.5, and an MCL inflation of 9.0 (13). Phylogenomics was performed using the Anvi’o (35). The pangenome was used to identify an optimal set of core genes to perform concatenated protein sequence phylogeny on, as described in the tutorial for the Anvi’o Pangenomics tutorial (https://merenlab.org/2016/11/08/pangenomics-v2/). The pangenomics analysis identified 348 single-copy genes encoded by all 244 genomes (i.e. single-copy core genes [SCGs]). Filtering by a minimum geometric homogeneity index of 1 left 270 core genes. Low functional homogeneity indices of these genes result in only 1-2 genes passing all filters, therefore the maximum functional homogeneity index was increased to 0.9925, leaving 12 optimal core genes on which to base the phylogenetic analysis of the 244 genomes (plus *Streptococcus sobrinus* as an outgroup). A list of the gene clusters in each genome, and the phylogenetic tree based on specific core genes were used as input for Coinfinder (14) to identify genes correlated to *muc* using the Bonferroni correction option. The output of Coinfinder was used by Cytoscape (36) to draw the correlation network in Figure 1.

### Generation of recombinant strains

#### (i) *mucG, mucH*, and *mucI* strains

B04Sm5 derivatives were constructed with the *mucG, mucH*, or *mucI* genes replaced by a spectinomycin resistance gene (specR), as previously described (8). Briefly, a 1010-bp fragment containing the spectinomycin resistance gene (specR) was amplified from pCAPB2 with primers spec_fwd and spec_rev. The left and right (∼500 bp each) flanking regions of *mucG, mucH*, or *mucI* were amplified from the genomic DNA of *S. mutans* B04Sm5 with the primer pairs of mucD_KO_L-fwd/mucD_KO_L-rev, mucD_KO_R-fwd/mucD_KO_R-rev, mucG_KO_L-fwd/mucG_KO_L-rev, mucG_KO_R-fwd/mucG_KO_R-rev, and mucI_KO_L-fwd/mucI_KO_L-rev, mucI_KO_R-fwd/mucI_KO_R-rev respectively. These three PCR products were assembled with a double digested pUC19 (PstI and EcoRI) using a NEBuilder HiFi DNA Assembly kit (New England Biolabs, USA), which resulted in the vectors pΔmucG, pΔmucH, and pΔmucI, respectively. These ligation products were transformed into *E. coli* NEB 5-alpha cells and positive clones were selected on LB agar medium containing 100 µg/ml spectinomycin. Vector clones were designated pΔmucG, pΔmucH, and pΔmucI, and verified by restriction analysis and sequencing. The disruption cassettes were amplified from constructs using primer pairs muc/G_KO_L-fwd/ mucH_KO_R-rev, mucH_KO_L-fwd/ mucG_KO_R-rev and mucI_KO_L-fwd/mucI_KO_R-rev, respectively. PCR products were digested by DpnI and then purified using the QIAquick PCR Purification Kit (Qiagen, USA). The disruption cassettes were transformed to *S. mutans* B04Sm5 by a previously reported protocol (37). Δ*mucG, ΔmucH*, and Δ*mucI* deletion mutants were selected by growth on BHI agar supplemented with 1 mg/ml spectinomycin, and confirmed by PCR and sequencing.

#### (ii) *mucG*-C, *mucH*-C, and *mucI-C* complement strains

Complement strains of Δ*mucG, ΔmucH*, and Δ*mucI* were generated using a previously described protocol (38). Briefly, single-copy genomic insertion of either *mucG, mucH*, or *mucI* into the *gtfA* locus of the cognate Δ*mucG*, Δ*mucH*, or Δ*mucI* strain was performed. Primers pmtaG-C erm F and pmtaG-C erm R were used to amplify the *mucG* open reading frame, as well as the 129 bp intergenic region upstream (between *mucG* and *mucF*) of *mucG*. Primers pmtaH-C erm F and pmtaH-C erm R were used to amplify the *mucH* open reading frame, as well as the 178 bp intergenic region upstream of *mucH* (between *mucH* and *mucI*). Primers pmtaI-C erm F and pmtaI-C erm R were used to amplify the *mucI* open reading frame, as well as the 178 bp intergenic region upstream of *mucI* (between *mucH* and *mucI*). The streptococcal integration vector pBGE was a gift from Jose Lemos and has been previously described (38). pBGE was linearized using XbaI and BsrG1, and the *mucG, mucH*, and *mucI* PCR products were ligated into pBGE using the Gibson Assembly Cloning Kit (New England Biolabs) to generate pmucG-C, pmucH-C, and pmucI-C respectively. These ligation products were transformed into *E. coli* NEB 5-alpha cells and positive clones were selected on LB agar medium containing 500 µg/ml erythromycin. The integrity of each construct was confirmed by sequencing. pmucG-C, pmucH-C, and pmucI-C were transformed into Δ*mucG* Δ*mucH*, and Δ*mucI* to generate the strains *mucG-C, mucH-C*, and *mucI*-C, respectively. Selection was performed on BHI agar containing 5 µg/ml erythromycin, and the integrity of both the deletion and complementation loci was confirmed by sequencing.

### HPLC

50 ml aliquots of BHI medium containing 1% glucose were inoculated with a loop of glycerol stock of B04Sm5, Δ*mucD*, Δ*mucF*, Δ*mucG*, Δ*mucH*, Δ*mucI, mucG-*C, *mucH*-C, or *mucI*-C and incubated at 37°C under 5% CO_2_ / 95% air. After 16 hours, 1 g of Amberlite XAD7-HP resin (Sigma Aldrich, Burlington, MA, USA) was added to the cultures. The cultures were then incubated for an additional 36 hours, after which the resin was recovered using a coffee filter. The resin was washed twice with 10ml of molecular grade water and extracted with 5 ml of ethyl acetate. The organic phase was decanted and evaporated, with the resulting pellet resuspended in 100 µl of methanol. Extracts were monitored at 280 nm during separation using an Agilent Technologies (Santa Clara, CA, USA) 1200 Series HPLC equipped with a Kinetex C18 100 Å LC column (5 µm, 150 × 2.1 mm)(Phenomenex, Inc., Torrance, CA, USA) as follows:

<u>Solvent A:</u> HPLC grade H_2_O : trifluoroacetic acid (TFA) (999 : 1, v/v)

<u>Solvent B:</u> HPLC grade acetonitrile (CH_3_CN) : trifluoroacetic acid (TFA) (999 : 1, v/v)

0-15 min: 30% solvent B

15-16 min: ramp 30%-100% solvent B

16-25 min: 100% solvent B

26-27 min: ramp 100%-30% solvent B

28-35 min: 30% solvent B

### Growth Curve

Ten microliters of overnight cultures of UA159, B04Sm5, Δ*mucD*, Δ*mucF*, Δ*mucG*, Δ*mucH, ΔmucI, mucG*-C, *mucH*-C, or *mucI*-C was added to 200 μl of BHI in a 96-well plate. Growth was monitored using a Tecan Infinite Nano. Optical density at 600 nm (OD600) was measured every hour for 20 h under 37 °C, with 5 s of shaking prior to each reading. Eight replicates of each strain were monitored.

### Deferred antagonism assay

The deferred antagonism assay was performed as previously described (8). Briefly, 8 µl of overnight cultures of UA159, B04Sm5 Δ*mucD*, Δ*mucF*, Δ*mucG*, Δ*mucH*, Δ*mucI, mucG*-C, *mucH*-C, or *mucI*-C was spotted onto BHI + 1% agar or BHI + 1% agar that was buffered to pH 7 with 1M KH_2_PO_4_/K_2_HPO_4_ pH 7.5, and incubated overnight at 37°C under 5%CO_2_/95% air. The following day, the plates were sterilized using the sterilization setting (90 s) in a GS Gene Linker UV Chamber (Bio-Rad, Inc.). 500 µl of overnight cultures of *S. sanguinis* SK36, *S. gordonii* ATCC10558, or *S. mitis* F0392 was added to 5 ml molten BHI + 0.75% agar that had been cooled to 40°C, and this was used to overlay the plates with the *S. mutans* colonies. The agar overlay was allowed to solidify at RT, and then the plates were incubated overnight at 37°C under 5%CO_2_/95% air. Zones of inhibition were measured the following day.

### RNA sequencing

B04Sm5, Δ*mucG*, and Δ*mucH* were grown in BHI at 37°C in 5% CO_2_ / 95% air to mid-log phase (OD_600_ ∼0.65 for B04Sm5 and Δ*mucH*; and 0.25 for Δ*mucG*, because of the growth defect phenotype). 1 ml aliquots of B04Sm5 and Δ*mucH* and 50 mL aliquots of Δ*mucG* were pelleted and frozen at -80° C. Total RNA was extracted using a RNeasy Power Microbiome Kit (Qiagen, Inc.) according to the manufacturer’s instructions. rRNA was depleted using a NEBNext rRNA Depletion Kit (Bacteria) (New England Biolabs) according to the manufacturer’s instructions. polyA tails were then added to the rRNA-depleted RNA using *E. coli* Poly(A) Polymerase (New England Biolabs). RNA was checked for quality at each step using a Qubit (Thermo Fisher Scientific) and a TapeStation (Agilent Technologies). RNA libraries were then constructed using the Direct cDNA Sequencing Kit (Oxford Nanopore Technologies) and sequenced on a GridION using a R9.4.1 Flow Cell (Oxford Nanopore Technologies). Basecalling was performed using Guppy v4.0.11/MinKNOW v20.06.09. Reads were mapped to the B04Sm5 genome using minimap2. The number of reads mapped to each gene was determined using featureCounts (39). Differential abundance between strains was determined using DeSeq2 (40) implemented using R (www.r-project.org). Genes with KEGG annotations were mapped on to the metabolic network using the KEGG color pathway tool (https://www.genome.jp/kegg/mapper/color.html). Minimap2 was used to map reads from the B04Sm5-, Δ*mucD-*, and Δ*mucF-*amended microbial communities published in (10) (available at NCVI; PRJNA773113) to the genome of B04Sm5. As above, the number of reads mapped to each gene was determined using featureCounts. Differential abundance between strains was determined using DeSeq2.

## Data availability

The raw mRNA sequencing reads used in this study have been deposited in the Sequence Read Archive (SRA) database and the BioProject accession number is PRJNA801007. The full *S. mutans* pangenome of 244 strains is available at https://github.com/jonbakerlab/Smutans-pangenome.

## Acknowledgements

The authors thank Karrie Goglin-Alemeida, Jelena Jablanovic, and Kara Riggsbee for performing the sequencing library preparation and sequencing, and Karen E. Nelson for helpful discussions. This research was supported by NIH/NIDCR F32-DE026947 (J.L.B.), K99-DE029228 (J.L.B.), R00-DE024534 (A.E.), and R21-DE028609 (A.E.).

## Figure Legends

**Figure S1: Organization of the *muc* BGC, and homology to the *rtc* BGC in *L. reuteri***. Diagram illustrating organization of the *muc* and *rtc* BGCs, the predicted functions of each gene product, and homology between cognate genes. Low homology between *mucF* and *rtcP*, and between the transcriptional regulators is indicated by red text and dashed lines. Adapted from Reference (8).

**Figure S2: *muc* pangenome correlation network**. Correlation network illustrating the *muc* genes (yellow diamonds) and genes they are correlated with, based on Coinfinder analysis. Nodes are other genes. Node color indicates presence of *muc* correlated genes in the shell (gray) or cloud (blue) pangenome. Node shape indicates gene clusters of interest: diamond = *muc*; chevron = hybrid NRPS/PKS; triangle = type-A2 lanthipepetide; circle = all other genes. Edges represent positive correlations only, and edge thickness indicates the p-value of the correlation.

**Figure S3: *S. mutans* phylogeny**. Phylogenetic tree of 244 *S. mutans* genomes based on the concatentated protein sequences of 12 core genes, as described in the Materials and Methods. The tree is annotated with 3 layers indicating the presence of *muc*, GC0000189 (type-A2 lanthipeptide which was correlated with muc in Figure 1D), and GC00001910 (hybrid NRPS/PKS which was correlated with *muc* in Figure 1D). Presence/absence bars in the *muc* layer are colored based on the position of the *muc* BGC of the cognate genome in the *muc* phylogenetic tree in Figure 1C. *S. mutans* genomes of interest are labeled (B04Sm5, UA159, UA140, NN20205, and 35).

